# Therapeutic Drugs Targeting 2019-nCoV Main Protease by High-Throughput Screening

**DOI:** 10.1101/2020.01.28.922922

**Authors:** Yan Li, Jinyong Zhang, Ning Wang, Haibo Li, Yun Shi, Gang Guo, Kaiyun Liu, Hao Zeng, Quanming Zou

**Author notes:** These authors contribute equally to this paper (Yan Li, Jinyong Zhang). Corresponding author: Hao Zeng, Quanming Zou.

## Abstract

2019 Novel Coronavirus (2019-nCoV) is a virus identified as the cause of the outbreak of pneumonia first detected in Wuhan, China. Investigations on the transmissibility, severity, and other features associated with this virus are ongoing. Currently, there is no vaccine or therapeutic antibody to prevent the infection, and more time is required to develop an effective immune strategy against the pathogen. In contrast, specific inhibitors targeting the key protease involved in replication and proliferation of the virus are the most effective means to alleviate the epidemic. The main protease of SARS-CoV is essential for the life cycle of the virus, which showed 96.1% of similarity with the main proteaseof 2019-nCoV, is considered to be an attractive target for drug development. In this study, we have identified 4 small molecular drugs with high binding capacity with SARS-CoV main protease by high-throughput screening based on the 8,000 clinical drug libraries, all these drugs have been widely used in clinical applications with guaranteed safety, which may serve as promising candidates to treat the infection of 2019-nCoV.

## 1. Introduction

Since December 2019, a series of pneumonia cases of unknown cause emerged in Wuhan, China. Genome sequencing of samples from these patients confirmed that the culprit of these infections was a beta-coronavirus that has never been reported before, which was later named as 2019-nCoV [1,2]. By Jan 28, 2020, more than 4500 patients were confirmed to be infected with 2019-nCoV, and nearly 100 patients were died from the infection. So far, the infection keeps spreading and more and more exported cases were confirmed in other provinces in China, and several other countries, including the United States, posing great pressure on public health security.

The discovery and clinical application of specific drugs against 2019-nCoV is an effective means to alleviate the current epidemic. However, there are no clinically effective drugs for this virus for the moment. Although 2019-nCoV is significantly different from SARS-CoV, which breakout in Beijing 17 years before [3], the sequence identity between themis as high as 79.5%. Further sequence alignment revealed that the similarity of the sequence of the main protease between 2019-nCoV and SARS-CoV is up to 96.1%. Previous study demonstrated that the main protease of SARS-CoV is essential for the life cycle of the virus, and is considered to be an attractive target for drug development [4]. Thus, this protein could be used as a homologous target to screening drugs that inhibiting the replication and proliferation of 2019-nCoV.

In order to screening the possible drug candidates that were able to prevent or cure the infection, high-throughput screening was performed based on the 8,000 clinical drug libraries using the online software Vina and SeeSAR, combined with our in-house automatic processing scripts and screening programs, and 4 small molecular drugs with high binding capacity with SARS-CoV main protease were identified. These drugs were widely used in clinical practice, so the safety was guaranteed. In view of the activity and safety of these wildly used drugs, it is feasible to conduct clinical exploratory treatment in special cases such as the outbreak of 2019-nCoV. Here we report the identify and structure of these molecules.

## 2. Material and Methods

### 2.1 Data resources

The CDS and proteins of Wuhan seafood market pneumonia virus (Accession: NC_045512.2) were obtained from NBCI database. The structures and sequences of SARS-CoV main protease were downloaded from PDB database. Nearly 8000 molecules were obtained from Drugbank [5], including the approval or experimental compounds and small molecules

### 2.2 Similarity search

Molecular similarity search was performed by using a strategy based on the similar sequences ofthe structure-revealed molecules. The NCBI-blast v2.9 was installed in local machine, then a local database was established by the molecular sequences of SARS-CoV main protease from PDB. The blastp was ran by default parameter.

### 2.3 Molecular docking

The AutoDock vina was obtained from website of scripps.edu [6]. The crystal structure of main protease monomer (PDB id: 5n5o) was used as target protein after removing the unrelated complex molecule by AutoDockTools. The receptor was prepared by removing water, adding hydrogen and computing charges. The binding coordinates were located by Grid box including the cave near N-terminal and the joint groove of dimer. Autodocking was performed by using multithread tasks by our in-house script, and potential molecules were screening out by a Perl program developed by us.

### 2.4 Protein-ligand interaction Analysis

The candidates harvested from docking were deeply analyzed for atom-based affinity contributions and physical-chemical properties by SeeSAR (version 9.2; BioSolveIT GmbH, Sankt Augustin, Germany, 2019, www.biosolveit.de/SeeSAR).

### 2.5 Structure Display

The 3D molecule images were displayed by PyMOL v2.3 [7].

### 2.6 Screening criteria

The affinities of Vina less than −7.7 kcal/mol were harvested initially. Then those molecules were removed such as toxin, experimental and unapproved ones, and those with strong side effects. Finally, the candidate drugs were selected with better affinities in Vina and SeeSAR.

## 3. Results

### 3.1 Homologous targets screening

We downloaded the sequences of a total of 103 SARS-CoV main proteases with different origins from PDB and built a local library for homology alignment. The result revealed that the similarity between the region at 3264 ∼ 3570 aa of 2019-nCoV ORF1 ab protein and SARS-CoV main protease 5n5o was up to 96.1% (Fig. 1). Thus, the main protease is highly conserved and main protease 5n5ocan be used as a homologous target for screening of possible drugs against 2019-nCoV.

**Fig. 1.**
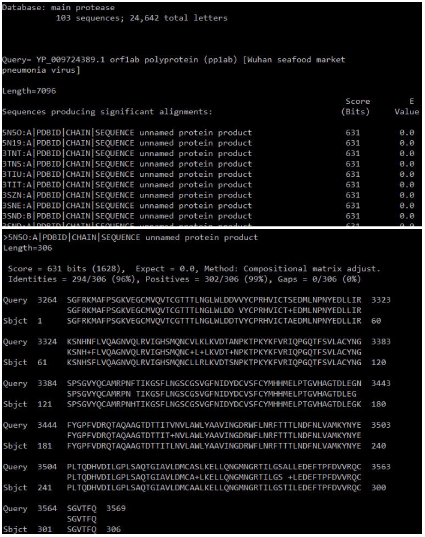
The blast results of ORF1ab of 2019-nCoV against SARS-CoV main proteases.

### 3.2 Binding sites of target protein detection

There were three binding pockets detected in the viral main protease by computation, as labeled in Fig.2C. In comparison to the dimer and monomer structure of the protein (Fig.2A and B), we could figure out that the pocket 1 was the binding site with natural substrate, pocket 2 was the joint groove between two monomers, and a small pocket 3 was located in the C terminal.

**Fig. 2.**
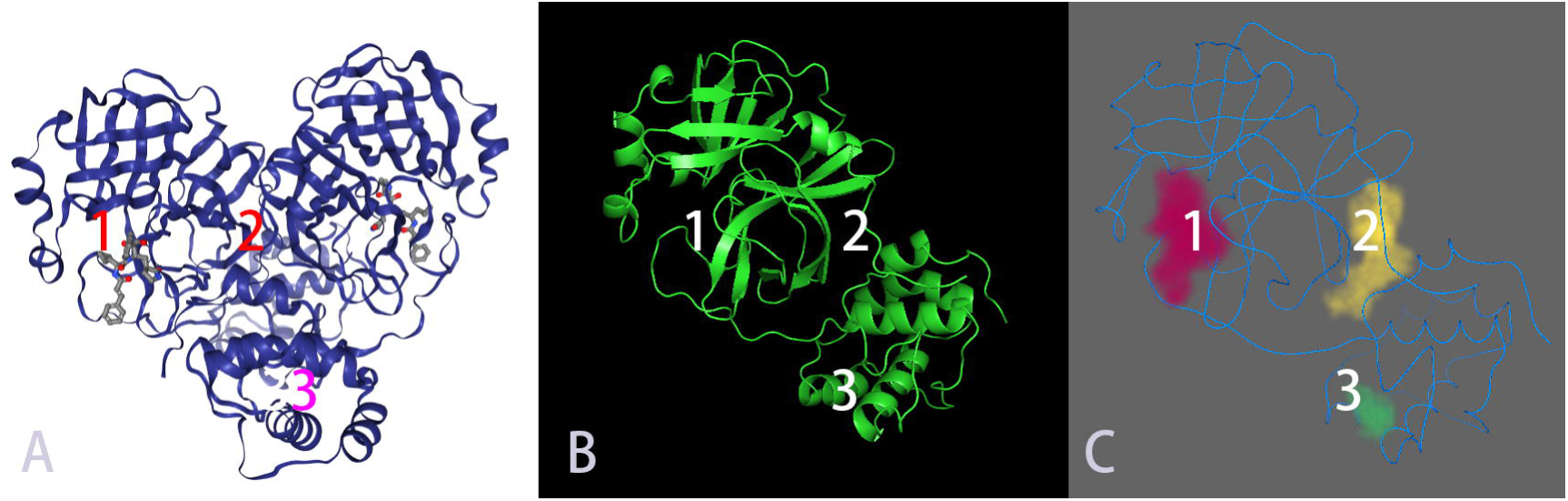
Binding pockets and functional sites of SARS-CoV main protease A: Dimer structure; B: Monomer structure; C: Detected pockets.

### 3.3 Candidates summary

At first, we harvested 690 molecules with possible binding capacity with SARS-CoV main protease, but most of them were dyes, toxins and antitumor drugs with strong side effects, about 50 molecules were left after excluding these molecules and other neurologic drugs, and then we selected marketable drugs from them for further kinetic and bio-chemical analysis. Finally, 4 molecules were identified, including No. 6651 molecule (Prulifloxacin), No. 6589 molecule (Bictegravir), No. 0097 molecule (Nelfinavir) and No. 6626 molecule (Tegobuvi).

### 3.4 The interaction between Prulifloxacin and viral protease

The binding energies of No. 6651 molecule (Prulifloxacin, Fig. 3A) to viral main protease were −8.2 Kcal/mol, −8.2 Kcal/mol, −7.9 kcal/mol at three binding sites, respectively (Fig. 3B), which were the cave adjacent to the N-terminal, the dimer joint groove and its back side. Moreover, at several docking poses, all of the estimated affinitiesof binding were between μM to mM. The evaluations of ligand-lipophilicity efficiency (LLE), torsion and clash were also satisfied.

**Fig.3.**
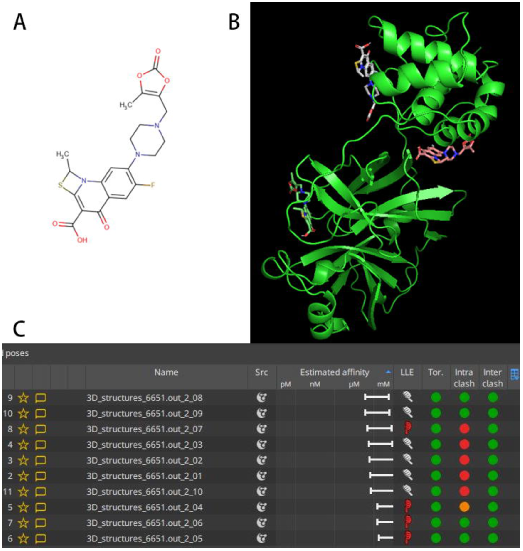
2D molecular structure of Prulifloxacin (A); Binding conformation of Prulifloxacin to viral protease (B); Physicochemical featureand affinity of Prulifloxacin to viral protease (C).

### 3.5 The interaction between Bictegravir and main protease

The binding sites between Bictegravir and the main protease was displayed at Fig. 4B, the binding energy between Bictegravir and major protease was −8.3 kcal/mol at the joint groove, but only −7.3 kcal/mol at the active site, and the estimated affinity is satisfied (Fig. 4C).

**Fig. 4.**
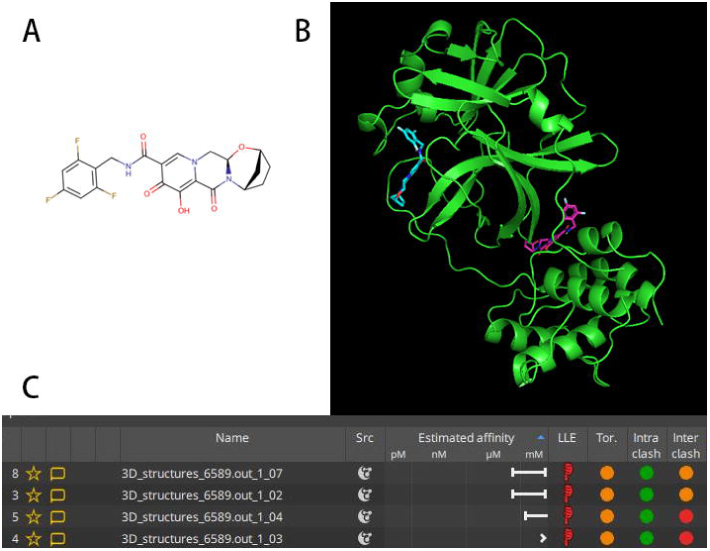
2D molecular structure of Bictegravir (A); Binding conformation of Bictegravir to viral protease (B); Physicochemical featureand affinity of Bictegravir to viral protease (C).

### 3.6 The interaction between Nlfinavir and main protease

The results showed that the binding site between Nelfinavir and the main protease was located at the joint groove (Fig. 5B). The binding energy between Nelfinavir and major protease was −8.6 kcal/mol, but the estimated affinity seemed not high enough (Fig. 5C).

**Fig. 5.**
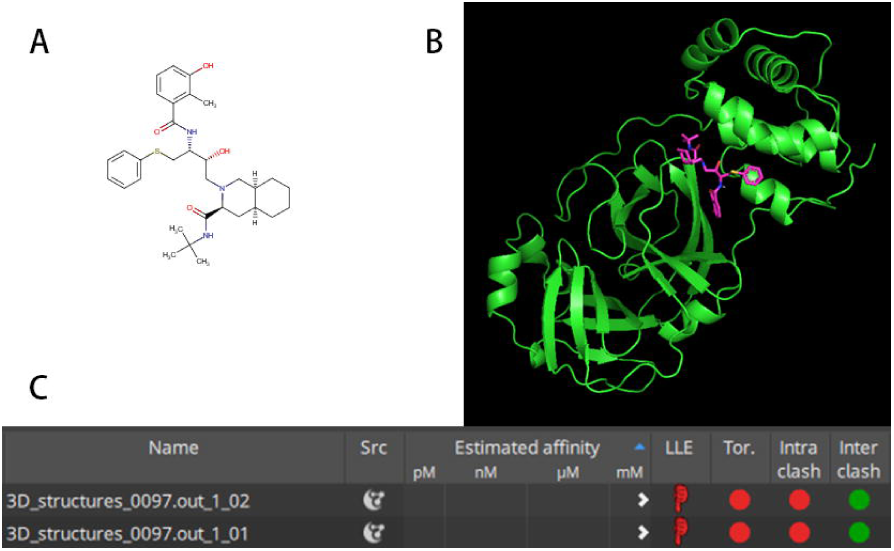
2D molecular structure of Nelfinavir (A); Binding conformation of Nelfinavir to viral protease (B); Physicochemical feature and affinity of Nelfinavir to viral protease (C).

### 3.7 The interaction between Tegobuvir and main protease

Tegobuvir was docked into the joint groove of main protein, the binding energy between them was −8.9 kcal/mol, and the estimated affinity is satisfied (Fig. 6C). The binding sites was displayed at Fig. 6B.

**Fig.6.**
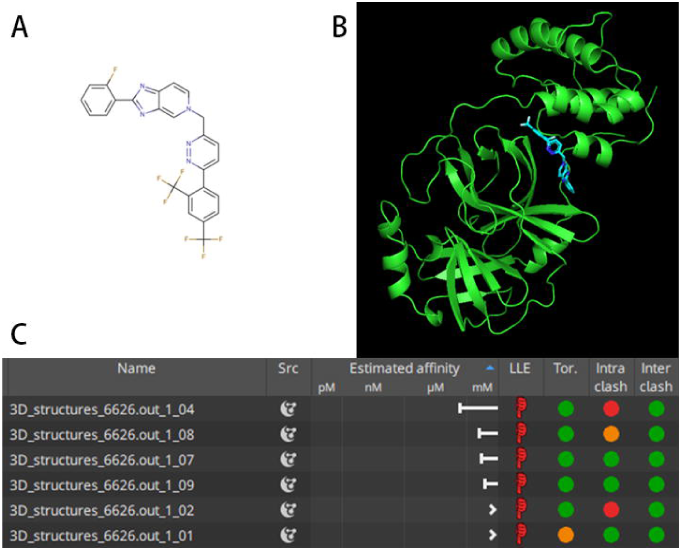
2D molecular structure of Tegobuvir (A); Binding conformation of Tegobuvir to viral protease (B); Physicochemical feature and affinity of Tegobuvir to viral protease (C)

## 4. Discussion

Although coronaviruses are under extensive mutagenesis, some key proteins, especially replication-related enzymes, are highly conserved [8]. Because the mutation in key proteins are often lethal to the virus [9], drugs that target conservative protease are usually capable of preventing the replication and proliferation of the virus and exhibit broad spectrum. Besides, they can reduce the risk of mutation mediated drug-resistance.

In the current study, based on the results from bioinformatics analysis, the structure of SARS-CoV 5n5o protein was selected as a homologous target for molecule screening. Then, in silico high throughput screening strategy and automatic pipeline have been established by using classic docking software and our in-house program, which greatly accelerates the screening process.

Among the four molecules identified in this study, Prulifloxacin is a chemotherapeutic fluoroquinolone antibiotic with broad-spectrum activity, it has been approved for the treatment of uncomplicated and complicated urinary tract infections, community-acquired respiratory tract infections in Italy and gastroenteritis, including infectious diarrheas, in Japan [10]. Unfortunately, Prulifloxacin is reported as a prodrug, which is rapidly metabolized to ulifloxacin *in vivo* [11]. But it can be used as a lead compound that provide guidance for our later structural modification and optimization to design more effective main protease inhibitors. Tegobuvir is a novel non-nucleoside inhibitor (NNI) of HCV RNA replication with demonstrated antiviral activity in patients with genotype 1 chronic HCV infection [12]. In addition, both of Nelfinavir and Bictegravir are anti-HIV drugs, of which Nelfinavir is a protease Inhibitors inhibit the cleavage of the polyprotein gag-pol [13], whereas Bictegravir is a new and potent HIV-1 integrase inhibitor, which is able to efficiently prevent HIV from multiplying and can reduce the amount of HIV in the body [14].

Our results clearly showed that all the four molecules showed reasonable binding conformations with the viral main protease. Among them, Prulifloxacin exhibited three binding sites with main protease, two of which were the same as Nelfinavir and Bictegravir. Further, molecule Prulifloxacin, Tegobuvir and Bictegravir are preferable to Nelfinaviras confirmed by affinity and physical-chemical properties analysis using SeeSAR. Based on the pockets’ functions of target protein, it suggested that these molecules should possess the abilities to block the active sites or interrupt the dimer formation of viral protein. Therefore, they may sever as promising candidates for drug repurpose and development against 2019-nCoV.

